# Sociality shapes the adaptive response of breeders to harsh thermal environments? An experiment in a burying beetle

**DOI:** 10.1101/2024.11.06.622234

**Authors:** Donghui Ma, Long Ma, Maaike A. Versteegh, Jan Komdeur

## Abstract

1. The rapid pace of environmental change seems to outstrip the evolutionary response rates of many species. Physiological, morphological, and behavioral plasticity alone may also not suffice to cope with environments’ drastic changes, potentially hindering individuals’ ability to adapt. Therefore, it has been proposed that social flexibility may be a powerful mechanism for fast adaptation, but empirical evidence is lacking.
2. We investigated the combined effects of manipulated social conditions and ambient temperature during the reproductive phase on parental care, reproductive success, and offspring performance of the burying beetle (*Nicrophorus vespilloides*), a facultative social breeding insect.
3. We conducted a mixed factorial design experiment, establishing four categories of social conditions during breeding (no-care, unifemale-care, biparenta-care, and multiple-care) alongside two thermal conditions (benign: 20°C and harsh: 23°C). Additionally, dispersed larvae produced from each of these breeding conditions were allocated to climate rooms maintained at either 20°C or 23°C for pupation. Data on brood size, brood mass, dispersed larval weight, larval development duration, body size and lifespan of newly eclosed beetles, and parental care were collected
4. Our results showed that (1) harsh temperature and the without-care condition (i.e., no-care compared to the other three social conditions) reduced brood size, brood mass, body size of newly eclosed beetles; (2) social and ambient thermal conditions had interactive effects on dispersed larval weight, larval development duration, and newly eclosed beetles’ lifespan; and (3) harsh temperature reduced both individual and total parental care, whereas sociality had opposing effects: individual care decreased, but total care increased from unifemale-care to biparental-care and multiple-care conditions.
5. Our findings shed light on the critical role of sociality in modulating the adaptive response of individuals to harsh thermal environments, one of the ecological problems exacerbated by ongoing global warming.

## 1 INTRODUCTION

Climate change, largely induced by global warming, currently has become one of the largest survival threats for animals and other organisms. Increasing evidence indicates that harsh and stressful thermal environments impact various aspects of reproduction, including sperm quality (e.g., Chevrier et al., 2019; Porcelli et al., 2017), fecundity (e.g., Pilakouta & Baillet, 2022), hatching success (Telemeco et al., 2009), as well as parental and offspring performance during breeding (e.g., Fragueira et al., 2021; Pilakouta et al., 2023; 2024; Vasudeva et al., 2021; Wiley & Ridley, 2016). These effects may have profound implications for species survival and persistence (e.g., Martinet et al., 2021; Rukke et al., 2018). Thus, it is essential to investigate the capacity and rate at which animals adapt to climate change (reviewed by Komdeur & Ma, 2021). Organisms possess the ability to adjust their morphology, physiology, and behaviour in response to a changing environment through evolutionary adaptation or phenotypic plasticity (reviewed by Komdeur & Ma, 2021). Organisms relying on evolutionary change for adaptation to environmental change require significant time scales for gene mutation and subsequent changes in allele frequencies to occur (e.g., Danchin et al., 2011; Grindstaff et al., 2003). In contrast, phenotypic plasticity enables animals to cope with the rapid pace of environmental changes much faster. Among various forms of phenotypic plasticity, behavioral plasticity stands out as one of the most important methods facilitating adaptation (e.g., Both et al., 2004; Chevin & Lande, 2015; Leimar, 2009; Nunney, 2016; Pigliucci, 2005). For example, birds may breed early in a warm year but later in a cold year to optimize reproduction (Visser & Both, 2005). In addition, sociality plays a crucial role in enabling animals to colonize novel environments (Cornwallis et al., 2017; Henriques & Osmond, 2020; Jarrett et al., 2017; Moss & While, 2021), but is often overlooked in its role of enhancing rather than limiting phenotypic plasticity in order to adapt to climate change (Komdeur & Ma, 2021). Therefore, it is important to explore whether and how sociality and phenotypic plasticity enhance species adaptation to challenging thermal across animal species.

Social behaviour among conspecific individuals encompasses a spectrum of cooperative and competitive interactions that occur during specific phases over their lifetime (e.g., reproduction) and are influenced by the status, condition, and/or genetic relatedness of the individuals involved (Ozgul et al., 2010; Paniw et al., 2019; Taborsky et al., 2021). Sociality serves as an index of the complexity of social structures within species, with highly social species typically displaying intricate social dynamics (e.g., Cornwallis et al., 2017; Hatchwell & Komdeur, 2000; Silk, 2007), whereas non-social (solitary) species exhibit simpler, singular behaviours and interactions that still reflect aspects of social dynamics (e.g., Makuya & Schradin, 2024). As John Donne (1624) famously noted, “no man is an island”; social behaviour is pervasive across species. For example, the widespread occurrence of social monogamy is common in bird species (Kempenaers, 2022), while eusocial insects such as ants and bees (order Hymenoptera) have demonstrated remarkable success in colonizing diverse habitats globally (Nowak et al., 2010; Ruxton et al., 2014). Social behaviour has been extensively examined in cooperative breeding species across a wide range of taxa, including insects, spiders, fishes, birds, and mammals (Field et al., 1998; Grinsted et al., 2014; Heg et al., 2004; Lukas & Clutton-Brock, 2012; Riehl, 2013). Recent theoretical and comparative studies have demonstrated that cooperative breeding initially evolved under stable and benign environments, but that cooperative breeders may have later outcompeted solitary breeders in regions with unpredictable, fluctuating, and harsh conditions(Branstetter et al., 2017; Gonzalez et al., 2013; Jetz & Rubenstein, 2011; Rubenstein & Lovette, 2007; Shen et al., 2017). However, direct evidence supporting social breeders outperform non-social breeders in challenging environments remains limited, specifically concerning whether sociality facilitates breeders to cope with harsh thermal conditions. Here, we investigate this question using burying beetles *N. vespilloides*.

Burying beetles are well known for their elaborate parental behaviour and flexibility in breeding systems (Potticary et al., 2024; Scott, 1998), which is largely driven by ecological (such as variations in elevation and temperature, Sun et al., 2014; resource availability, Wang et al., 2022) and social dynamics (e.g., intra- and interspecific competition, Komdeur et al., 2013; Tsai et al., 2020). Upon locating a suitable resource (i.e. small vertebrate carcass) for breeding, both male and female beetles engage in indirect parental care by burying the carcass into the soil, removing the fur or feather, shaping it into a ball, and coating it with a mixture of oral and anal secretions (e.g., Shukla et al., 2018). Also, these beetles jointly defend the carcass against both intra- and interspecific rivals (e.g., Eggert & Müller, 1989; Müller et al., 1990; Trumbo, 2022). When larvae hatch, beetle parents provide direct care towards their developing larvae, such as offspring provisioning, and show considerable flexibility in parental coordination (e.g., Smiseth, 2005; Scott, 1998). Males and females often collaborate in raising their young (biparental care), but sometimes one parent can take care of offspring alone (i.e., uni-female or uni-male care) (Jarrett et al., 2017; Moss & Moore, 2021; Wang et al., 2021). Despite the positive impact of parental care on offspring development and fitness, in the absence of parents, larvae can normally develop and survive through self-feeding on the carcass in some burying beetle species (Attisano & Kilner, 2015; Jarrett et al., 2017). In other cases, multiple individuals of different sexes reproduce collectively by preparing and sharing the same single carcass, and they may take care of all offspring in the communal brood, known as co-breeding (Komdeur et al., 2013; Ma et al., 2022; Richardson & Smiseth, 2020). It has been reported that changes in environmental temperature affect reproductive success and offspring performance in burying beetles. Breeders appear to adjust their parental care in response to these thermal changes (e.g. *N. nepalensis*, Malik et al., 2024; *N. orbicollis*, Moss & Moore, 2021; *N. vespilloides*, Pilakouta et al., 2023, 2024). Their varied social structures during breeding (including no care, uniparental care, biparental care, and multiple caregivers) make burying beetles an ideal model species for exploring whether and how sociality enables animals to adapt to changing climatic conditions.

In this study, we address three key questions: (1) the impacts of harsh thermal conditions on reproductive success and offspring performance; (2) whether different social conditions can show distinct capacities of buffering reproductive costs associated with temperature changes; and (3) whether and how breeders can adjust their parenting behaviours and coordination in care in response to harsh thermal conditions. To explore these questions, we employed a mixed factorial design experiment to investigate the combined effects of social conditions and temperatures on reproductive success, offspring performance, and parental behaviour during breeding. We established four categories of social conditions -- no care, unifemale care, biparental care, and multiple care by manipulating the number and sex of adults. These beetles were then exposed to two temperature regimes (20 °C and 23 °C). We focused on unifemale care, as it is more common than unimale care in burying beetles (Bartlett, 1987; Scott, 1998).We defined 20°C as a benign thermal condition since the beetles have been acclimated to this temperature for several generations and exhibit successful breeding. In contrast, we have chosen a mild temperature increase to 23°C as harsh thermal condition based on previous studies (Moss & Moore, 2021; Pilakouta et al., 2023), and this temperature did not cause severe reproductive failures, compared as our pilot experiment showing that 25 °C led to no successful breeding (0/120 broods). We assessed a suite of proxies for reproductive success and offspring performance such as brood size, brood mass, average larval mass, and duration of larval development. We assessed parental care by measuring frequency spent on the carcass by parents. To further investigate the effects of thermal conditions during pupation on offspring performance and fitness, all dispersed larvae from different social conditions and thermal conditions were allocated to either 20°C or 23°C for pupation, and average size of newly-eclosed beetles and their lifespan were measured.

In accordance with previous studies showing that harsh thermal conditions adversely affect reproduction and offspring fitness in burying beetles (e.g., Pilakouta et al., 2023, 2024), we expect such adverse impacts for all social conditions. However, we expect that the different social conditions have varying capacities to buffer against these harsh thermal conditions, and such buffering capacity of social conditions may result from adjustments in parental behavior and coordination among breeders. The presence of breeding mates and other group members (i.e., co-breeders) is found to enhance fitness benefits in reproduction, such as increased parental care and improved resources defence (Eggert et al., 1998; Liu et al., 2020; Ma et al., 2022; Tsai et al., 2020). However, such benefits may be largely hidden or diminished in benign environments (García-Ruiz et al., 2022; Rubenstein & Lovette, 2007), for example, comparable reproductive success has been found between bi-parental and uniparental care (Wang et al., 2021). We predict that under harsh thermal conditions, biparental and multiple-care conditions will outperform unifemale-care in reproductive success and offspring performance. Additionally, we anticipate that individuals in biparental and multiple-care conditions will adjust their parental behaviour and demonstrate greater coordination in parental care between partners in response to harsh thermal conditions, compared to those in the unifemale-care condition. We expect higher sociality to buffer reproductive costs in harsh environments, likely through rapid adjustments in parental behavior and enhanced coordination in care. Our study may reveal whether animals enhance their adaptation to stressful thermal conditions by adjusting social behaviors. Understanding the role of sociality in adapting to harsh environments will contribute to a broader comprehension of its function as an adaptive mechanism in response to environmental change.

## 2 METHODS

### 2.1 Beetles husbandry

All adult beetles used for this study were fifth-generation individuals derived from an outbred laboratory population at the University of Groningen, The Netherlands. This laboratory population originated from wild-caught beetles from De Vosbergen (53° 08’ N, 06° 35’ E), The Netherlands, using pitfall traps baited with thawed mice (bought from a pet store), during the reproductive seasons from June to July in 2022. The adults were brought to the lab, and were allowed to reproduce over the period August 2022 until April 2023. All adults and larvae were maintained at 20°C with a 16 h: 8 h light to dark cycle. Groups of up to six same-sex adult beetles from the same parents were kept in a clear rearing plastic box (10cm L × 6cm W × 8 cm H) with moist peat. Throughout the study, non-breeding beetles were fed twice a week with 2∼3 beheaded mealworms per beetle each time.

### 2.2 Experimental design and procedures

We experimentally conducted a 4 × 2 factorial design, where we established four social conditions during breeding. (i.e., no-care, unifemale-care, biparental-care, and multiple-care conditions) at two ambient breeding temperature conditions (20°C or 23°C) (Fig.1; for sample sizes see Supplementary Information S1).

**Figure 1.**
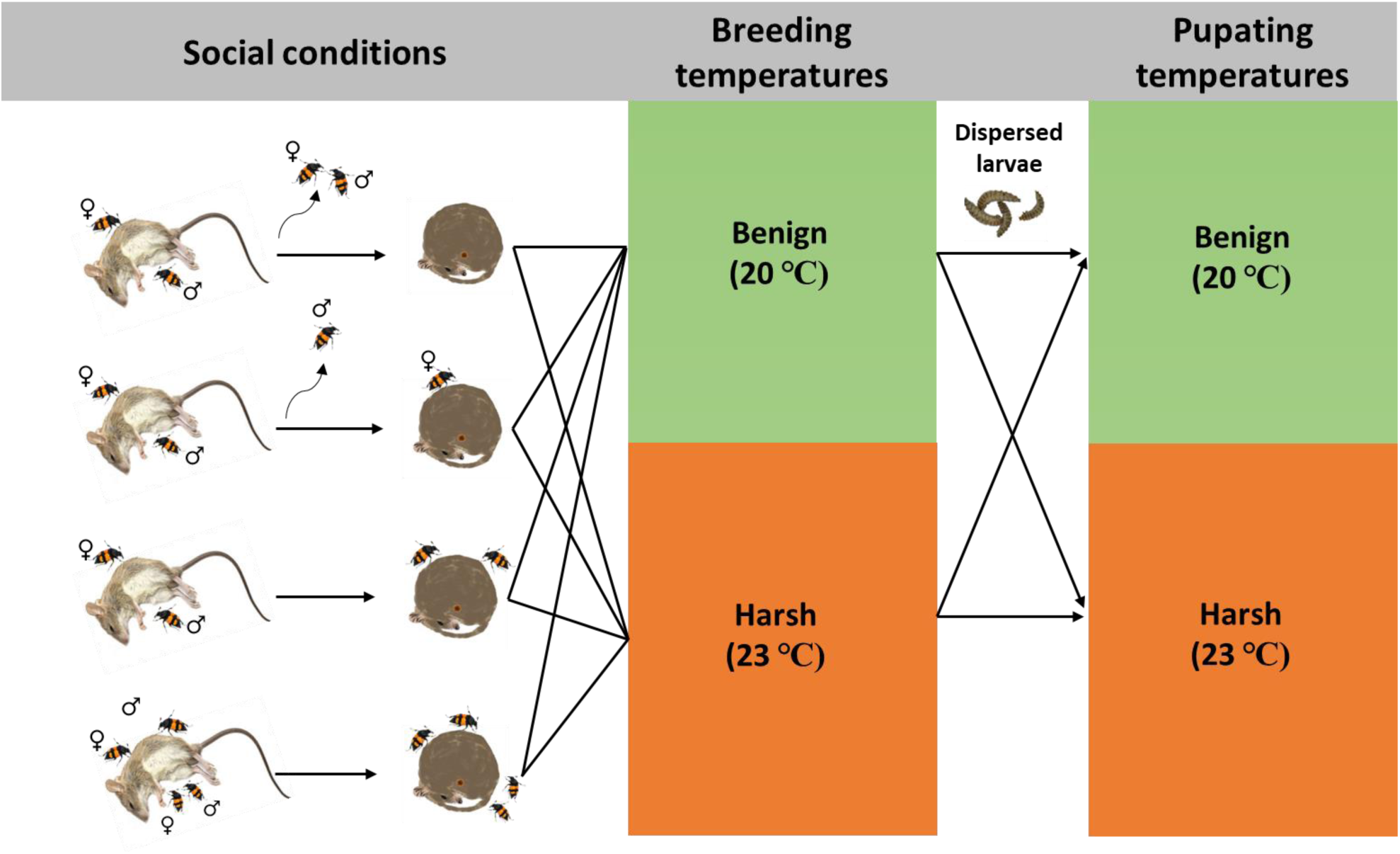
Experimental design of the study (not drawn to scale). The left section depicts four social conditions arranged vertically: no-care, unifemale-care, biparental-care, and multiple-care. The middle section shows two climate rooms set at 20°C and 23°C for breeding events. The right section illustrates the transfer of dispersed larvae to the 20°C and 23°C climate rooms for pupation.

Virgin, sexually mature beetles reared in a 20°C climate room, aged two weeks post-eclosion, were selected and their body size (pronotum width) was measured using an electric caliper (accuracy 0.01mm). Similarly sized (Supplementary Information S1), unrelated individuals (i.e., they nor their parents were siblings) of the opposite sex were paired and placed together in a plastic breeding container (23cm L × 19cm W × 12.5 cm H) filled with moist soil ranging from 3 to 5 cm in depth. For each breeding event, a newly thawed dead mouse, weighed (Supplementary Information S1) using an electronic balance (accuracy 0.01g), was provided to initiate breeding. All adults were individually marked by making small holes in the elytra with a 00 insect pin, facilitating the observers’ recognition of beetles. During the entire period of breeding (from the onset of the experiment until larval dispersal from the carcass), beetles and larvae activity were checked three times daily with intervals of 5 hours (08:00-09:30 am, 14:30-16:00, 21:00-22:30). During each check, the soil covering the carcass was removed gently, and the presence or absence of individual marked parents on or in the carcass was recorded (Ma et al., 2022; Wang et al., 2021, 2022). Upon egg hatching (i.e., the first instar larvae appear on the carcass), male and female parents were removed from the breeding containers to create the no-care treatment approximately 3 ± 1 days after initiating breeding, which ensures that the pair has mated and the carcass has been prepared sufficiently. To create the ‘unifemale-care’ condition, the males were removed when the first larva hatched. To create the ‘biparental-care’ condition, one pair of beetles was allowed to breed until larvae dispersed without any removal manipulation. To create the ‘multiple-care’ condition, two pairs of beetles were used for breeding on a single carcass, and all beetles were allowed to be present until larvae dispersed. Given the positive correlation between individual body size and competitive ability in burying beetles (e.g., Beeler et al., 2002; Otronen, 1988), the two pairs differed in body size (overall variation between dominant and subordinate individuals is 9.35%, Supplementary Information S1), to create a dominance hierarchy (Ma et al., 2022). Parental care behaviour was measured and defined as preparing and preserving the carcass, and provisioning larvae on the carcass (e.g., Eggert et al., 2008; Lee et al., 2014; Smiseth et al., 2005). As measures of parental care, we used the proportion of times that each individual (i.e. individual care) and the sum of all individuals (group care) in the box was present on the carcass throughout the entire reproductive event (e.g., Ma et al., 2022; Pilakouta et al., 2015; Wang et al., 2021, 2022). At larval dispersal (i.e., most of the third instar larvae leave from the carcass), for each brood we recorded the number of larvae (i.e., brood size) and weighed all larvae individually using an analytical balance (accuracy to 0.0001g) to obtain brood mass and average larval mass.

Finally, we examined whether larvae raised at different social conditions and temperatures showed variation in responding to benign and harsh conditions while pupating. Therefore, just before pupation (after about 3 weeks after dispersal) all dispersed larvae were transferred into containers with moist peat (one brood per container; volume of peat varied based on larval number). All containers were then allocated to two different temperatures (20℃ and 23℃). For pupation, half the larvae remained at the same temperature and half were allocated to the other temperature regime, leading to four different developmental conditions: 20℃-20 ℃ (N = 106), 20℃-23℃ (N = 53, see Supplementary Information S2, 23℃-20℃ (N = 54), 23℃-23℃ (N = 53). After eclosion, we counted and sexed the newly emerged beetles from each brood. Individual body size (pronotum width; 0.001 mm) was measured using ImageJ (version 1.53) (Bladon et al., 2020) based on images of each adult. We then randomly (i.e., excluding those that are too small or too large in body size) assigned four newly eclosed beetle adults (2 males and 2 females) per pupation container, with between 28 and 34 individuals per treatment group (Supplementary Information S3). These beetles were housed individually, fed beheaded mealworms three times weekly, and maintained at 20°C with a 16 h:8 h light cycle until death to record their lifespan.

We evaluated reproductive success by measuring brood size and brood mass. Furthermore, we calculated the average weight of dispersed larvae per brood to examine the effect of breeding temperature and the average body size of newly eclosed beetles per brood to examine the effects of breeding and pupation temperature on offspring performance. We recorded the timing of egg hatching and larvae dispersal in days, enabling the calculation of the duration of the larval developmental stages (i.e., the phase from the first egg hatching until the third instar larvae disperse).

### 2.3 Data analyses

We performed linear regression models for the two reproductive outcome traits, brood size and brood mass with explanatory variables social condition (*S*), ambient breeding temperature (*T_b_*) and their interaction (*S × T_b_*), and the covariate carcass mass. We conducted similar linear regression models for the duration of larval development. Secondly, linear regression models were employed on average larval mass and the size of newly eclosed beetles, using the same explanatory variables as above, and adding pupation temperature (*T_p_*) and the interactions (*S × T_p_*) for the models of average body size of newly eclosed beetles. The third variable used as a proxy for offspring performance is lifespan, which directly indicates individual survivability and reproductive potential. This was analyzed using the Cox proportional hazards model (*coxph* within the *survival* package, Terry & Patricia, 2000). The explanatory variables encompassed the same variables as were used in the model on average size of newly eclosed beetles. Finally, we conducted a binomial generalized linear mixed model (GLMM) for individual and total parental care with explanatory variables social condition, ambient breeding temperature and their interaction (*S × T_b_*), the covariate carcass mass, and the random factor breeding containers ID. We did *post hoc* testing for those models that exhibit significance in interactions or social conditions using *emmeans* from the *emmeans* package (Length, 2023). These data analyses were conducted in R 4.3.3 (R Core Team, 2024), and figures were generated using the *ggplot2* package (Wickham, 2016). The level of statistical significance for all tests was *P* = 0.05.

## 3 RESULTS

### 3.1 Effects of ambient thermal and social conditions on reproductive success and larval development

#### 3.1.1 Reproductive success: brood size and brood mass

Both ambient thermal and social conditions had significant effects on brood size and brood mass at larval dispersal (thermal condition: *F* = 10.52, *P* = 0.001 and *F* = 19.73, *P* < 0.001, respectively; social condition: *F* = 2.79, *P* = 0.04 and *F* = 16.36, *P* < 0.001, respectively), while their interaction did not (*S* × *T_b_*: *F* = 0.59, *P* = 0.62 and *F* = 0.91, *P* = 0.44, respectively). Compared to the benign temperature, in the harsh temperature smaller and lighter broods were produced. Compared to the no-care condition, larger brood size was only found in unifemale-care condition (Estimate = 3.75, SE = 1.35, *t* = 2.78, *P* = 0.03), but not in the biparental- and multiple-care conditions (Estimate = 2.65, SE = 1. 34, *t* = 1.98, *P* = 0.20 and Estimate = 1.42, SE = 1. 47, *t* = 0.97, *P* = 0.77, respectively, Fig.2a), whereas heavier broods were found in all these three social conditions (Estimate = 1.25, SE = 0.20, *t* = 6.28, *P* < 0.001; Estimate = 1.14, SE = 0.20, *t* = 5.8, *P* < 0.001; Estimate = 0.63, SE = 0.22, *t* = 2.94, *P* = 0.02, respectively, Fig.2b). Furthermore, broods that received care had a similar brood size (*t* = 1.54, *P* > 0.41, Fig.2a), whereas broods in multi-care condition were lighter than in the unifemale-care condition (Estimate = 0.61, SE = 0.22, *t* = 2.76, *P* = 0.03, Fig.2b).

**Figure 2.**
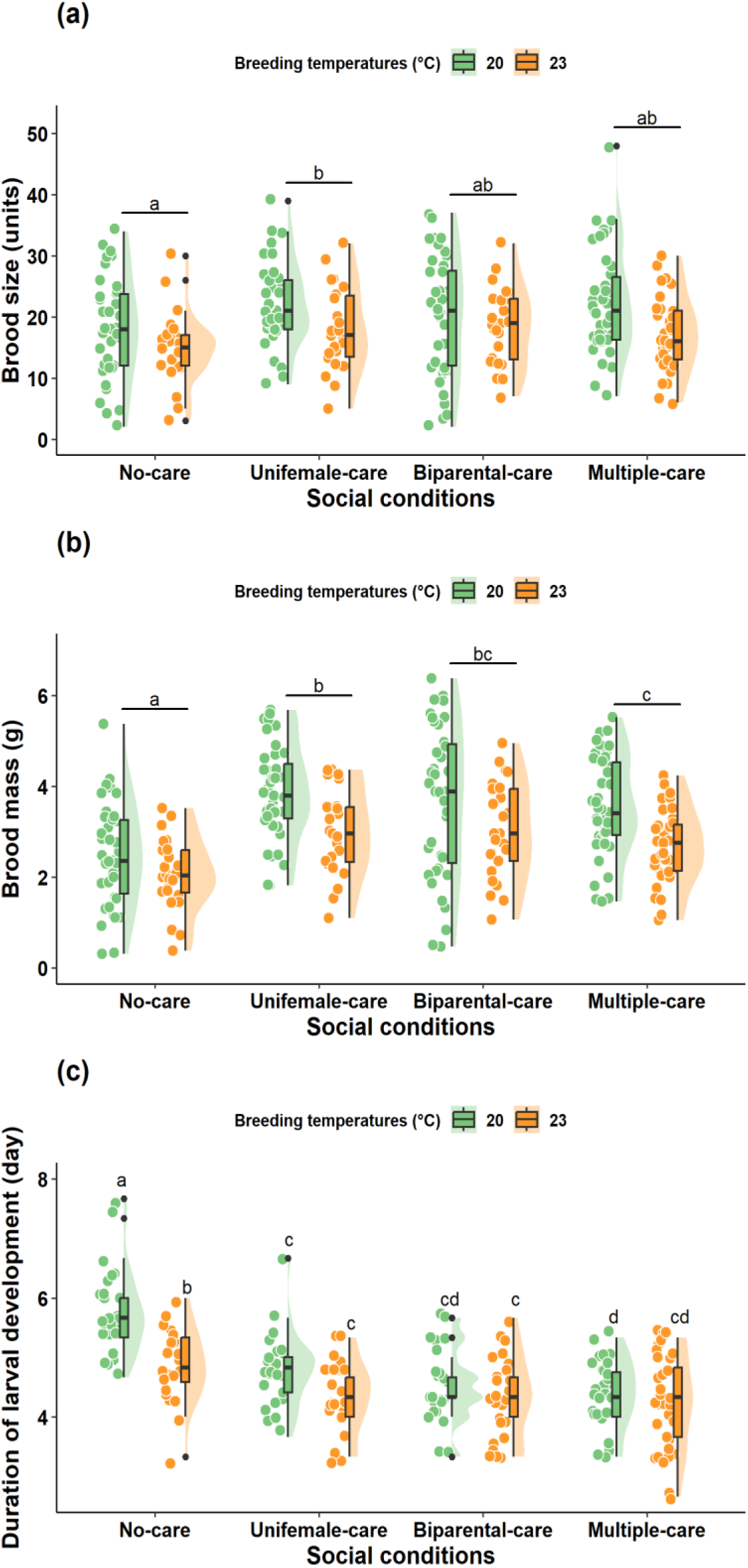
Effect of social and ambient thermal breeding conditions on (a) brood size, (b) brood mass, and (c) duration of larval development in burying beetles. Significant differences are indicated by letters (comparing only differences across social conditions in (a) and (b)). Coloured dots represent the sample values, and black dots are the outliers. The distribution of variables is drawn at the right of the boxes.

#### 3.1.2 Duration of larval development

We found a significantly interactive effect of thermal and social conditions on the duration of larval development (*S × T_b_*: *F* = 3.15, *P* = 0.03). In the no-care condition, the harsh thermal condition significantly shortened the duration of larval development compared to the benign thermal condition (Estimate = 0.73, SE = 0.2, *t* = 3.75, *P* < 0.001), whereas such an effect was not observed within any of the other three social conditions (*t* < 1.62, *P* > 0.11, Fig.2c). In both thermal conditions, the no-care condition showed the longest larval development compared to the other three conditions (*t* > 2.84, *P* < 0.03, Fig.2c). In the benign temperature, the multiple-care condition exhibited a shorter duration of larval development than unifemale-care condition (Estimate = 0.56, SE = 0.19, *t* = 2.97, *P* = 0.017), while such a difference was not found in the harsh temperature (Fig.2c).

### 3.2 Effects of ambient thermal conditions during breeding and pupation, and social conditions on offspring performance

#### 3.2.1 Average weight of dispersed larvae

The interaction between ambient temperature and social conditions significantly affected the average weight of offspring at larval dispersal (*S × T_b_*: *F* = 2.97, *P* = 0.03). Interestingly, only in the biparental-care condition, a decreased average weight of dispersed larvae was observed in the harsh ambient condition compared to the benign ambient condition (Estimate = 0.022, SE = 0.006, *t* = 3.96, *P* < 0.001). Furthermore, the lightest dispersed larvae were grown in the no-care condition, irrespective of temperature (*t* > 3.81, *P* < 0.001), while under benign breeding temperature, dispersed larvae growing in the biparental-care condition were heavier than those in the multiple-care condition (Estimate = 0.022, SE = 0.005, *t* = 4.63, *P* < 0.001) (Fig.3a).

**Figure 3.**
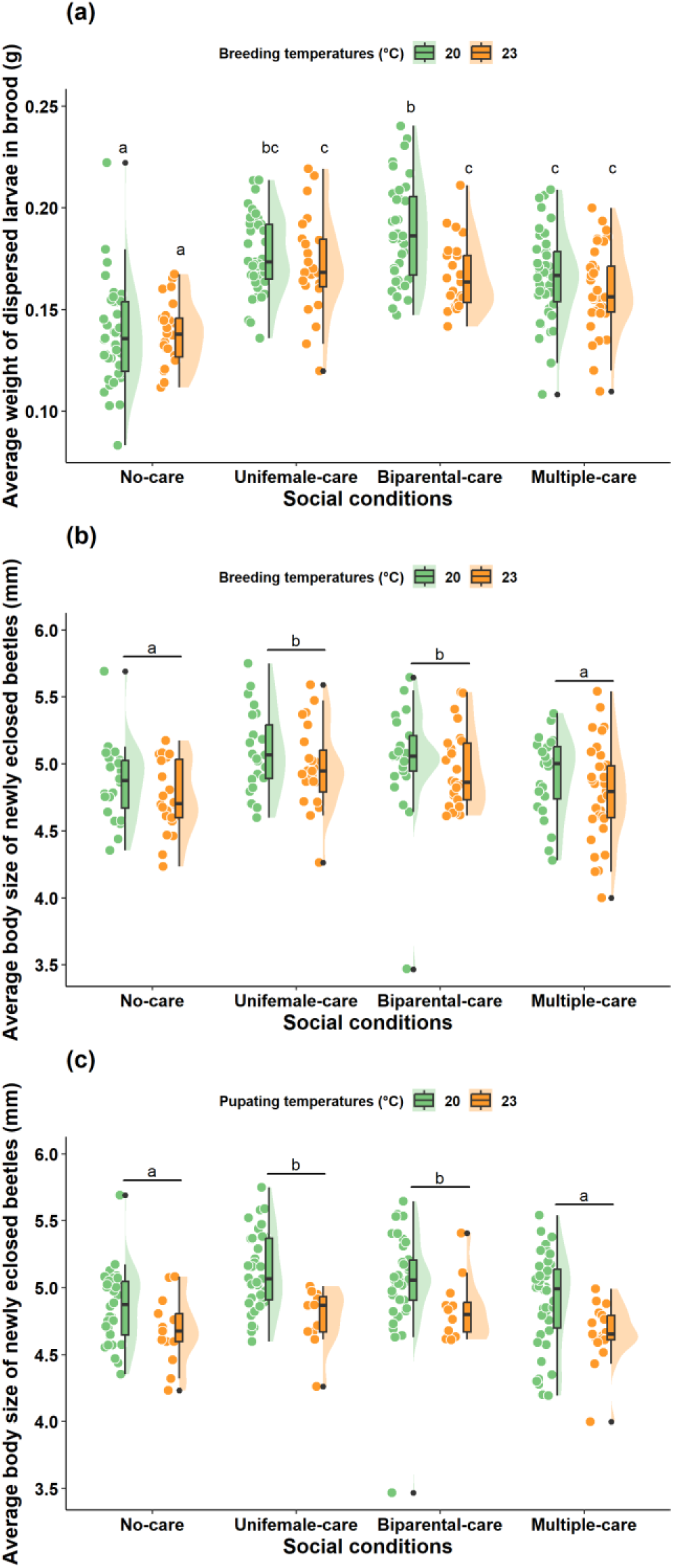
(a) Effect of ambient breeding temperature and social environment on the average weight of dispersed larvae, (b) the average body size of newly eclosed beetles, and (c) effect of ambient pupating temperature and social environment on the average body size of newly eclosed beetles. Significant differences are indicated by letters (comparing only differences across social conditions in (b) and (c)). Colo ured dots represent the sample values, and black dots are the outliers. The distribution is drawn on the right of the boxes.

#### 3.2.2 Average body size of newly eclosed beetles

The average body size of newly eclosed beetles was significantly influenced by ambient temperature during pupation (*F* = 7.46, *P* = 0.007) and social conditions (*F* = 8.51, *P* < 0.001, Fig.3c), but not by ambient temperature during breeding (*F* = 2.20, *P* = 0.14, Fig.3b), or either of the interactions (*S × T_b_* and *S × T_p_* : *F* < 0.43, df = 3, *P* > 0.73). Smaller newly eclosed beetles were produced in the harsh pupation temperature compared to the benign pupation temperature. Compared to the no-care and the multiple-care conditions, larger beetles were found in the unifemale-care (Estimate = 0.21, SE = 0.06, *t* = 3.35, *P* = 0.006; Estimate = 0.30, SE = 0.07, *t* = 4.15, *P* < 0.001 respectively) and the biparental-care conditions (Estimate = 0.18, SE = 0.06, *t* = 2.91, *P* = 0.02; Estimate = 0.27, SE = 0.07, *t* = 3.7, *P* = 0.002, respectively) (Figs 3b, c).

#### 3.2.3 Offspring lifespan

The results of Cox proportional hazards models showed that neither ambient breeding temperatures (*χ^2^* = 0.26, *P* = 0.61, Fig.4a) nor the interaction of ambient breeding temperature and social conditions (*S × T_b_*: *χ^2^* = 3.77, *P* = 0.29) did affect offspring lifespan. However, the interaction between ambient pupation temperature and social condition was of significant effect (*S × T_p_*: *χ^2^* = 8.78, *P* = 0.03; Figs 4b, c). In comparison to the benign pupation temperature, offspring beetles from the harsh pupation temperature experienced higher mortality rates only under biparental care (Estimate = 0.84, SE = 0.19, *z* = 4.53, *P* < 0.001), with no significant effects observed under the no-care, unifemale-care, or multiple-care conditions (*z* < 1.75, *P* > 0.08). Additionally, when pupating at the benign temperature, newly eclosed beetles in the multiple-care condition experienced higher mortality rates than those in the no-care (Estimate = 0.48, SE = 0.18, *z* = 2.62, *P* = 0.04) and the biparental-care conditions (Estimate = 0.53, SE = 0.18, *z* = 2.87, *P* = 0.02). In contrast, when pupating at the harsh temperature, newly eclosed beetles in the biparental-care condition faced a higher risk of mortality than those in the no-care condition (Estimate = 0.476, SE = 0.18, *z* = 2.59, *P* = 0.048) (Figs 4b, c).

**Figure 4.**
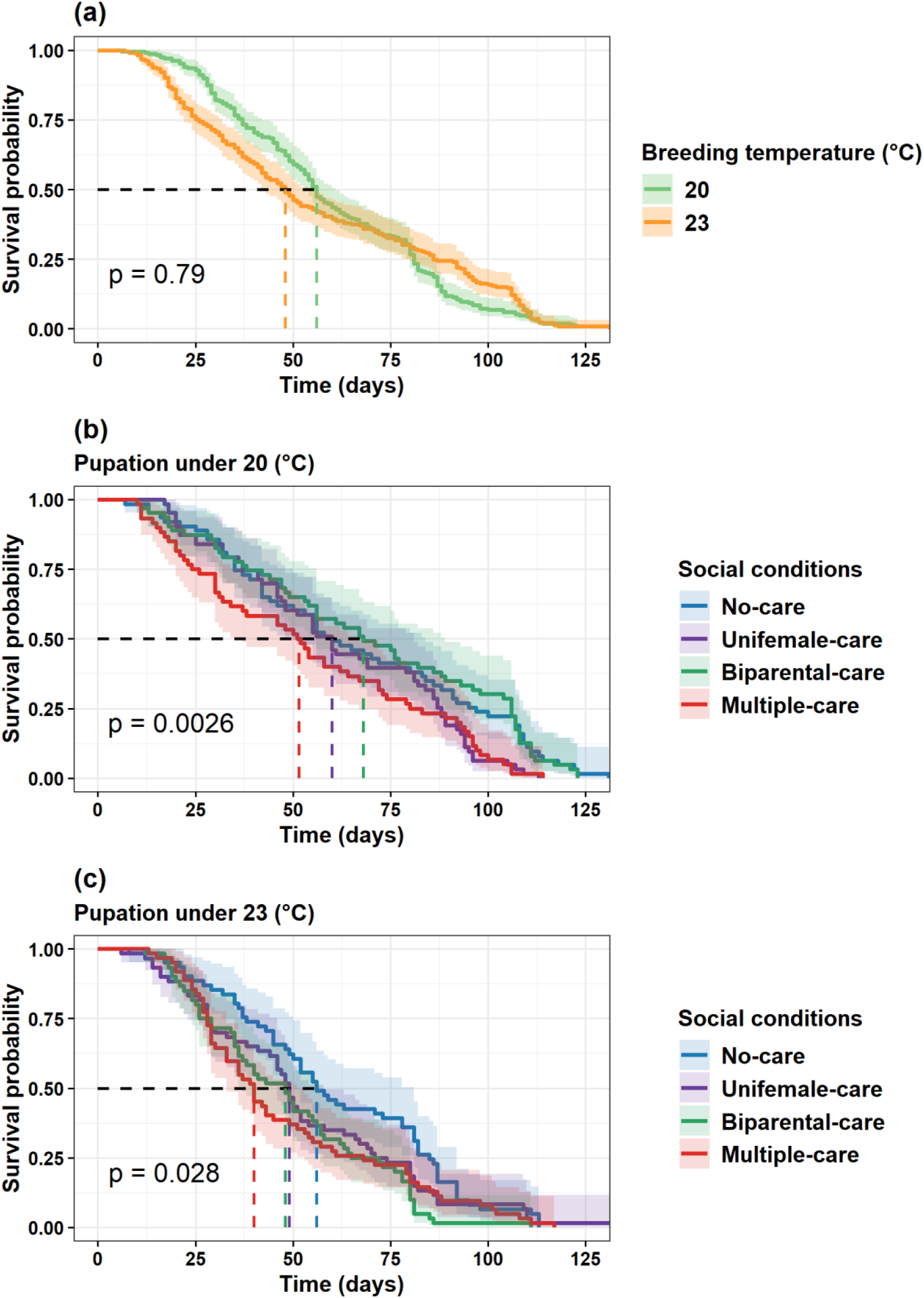
The effect of (a) breeding temperatures, and the effect of social environment in (b) benign, and (c) harsh pupation temperature on the lifespan of offspring burying beetles.

### 3.3 Effects of ambient thermal and social conditions on parental care

We found significant effects of both ambient thermal and social conditions on individual and total parental care (thermal condition: χ² = 6.82, *P* = 0.009 and *χ*² = 4.26, *P* = 0.04, respectively; social condition: *χ*² = 32.12, *P* < 0.001 and *χ*² = 178.28, *P* < 0.001, respectively), but no effect of their interaction (*S × T_b_*: *χ*² < 0.37, *P* > 0.83). Individuals reduced their parental care in response to the harsh thermal condition, providing lower total parental care compared to the benign condition. Individual parental care was lower in the multiple-care and biparental-care conditions compared to the unifemale-care condition (*z* > 4.87, *P* < 0.001, Figure 5a). However, total parental care varied significantly across social contexts, with the highest care observed in the multiple-care condition, followed by the biparental-care, and the unifemale-care conditions (pairwise comparisons: *z* > 5.39, *P* < 0.001, Fig. 5b).

**Figure 5.**
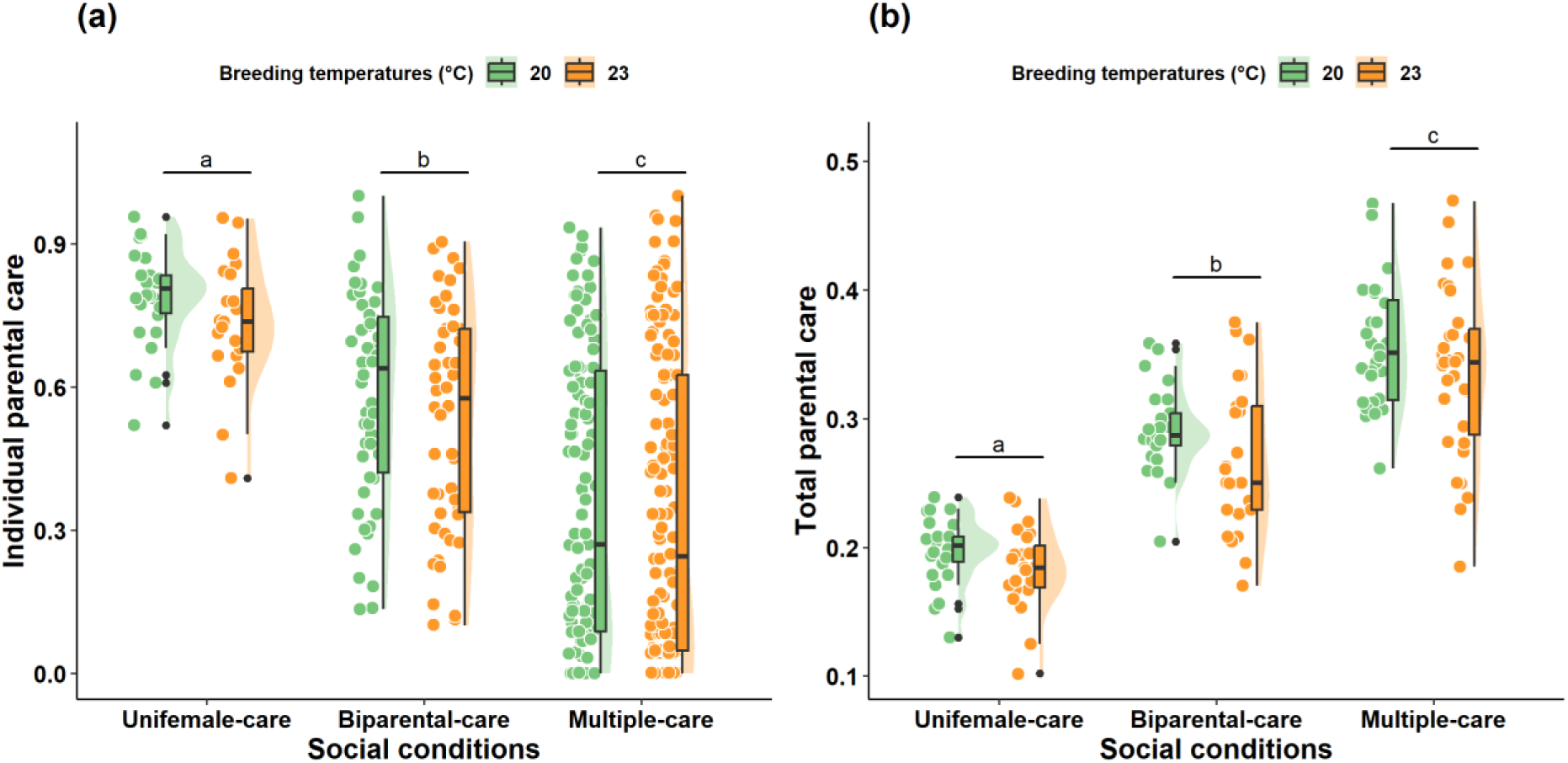
(a) The effect of social conditions (excluding the No-care condition) on (a) total parental care, (b) individual parental care. Significant differences are indicated by letters (comparing only differences across social conditions). Colored points represent the sample values, and black points are the outliers. The distribution is drawn on the right of the boxes.

## 4 DISCUSSION

Sociality may facilitate the adaptation of animals to challenging physical environments (e.g., Jarrett et al., 2017; Komdeur & Ma, 2021; Moss & While, 2021). Here we test this hypothesis by experimental manipulation. Firstly, as predicted, we found that harsh ambient temperatures during breeding increased reproductive costs for breeders and negatively affected offspring performances. Secondly, we found that compared to the no-care condition, the other three social conditions exhibited improved reproductive success and offspring performance, with unifemale-care and biparental-care having a stronger effect than multiple-care. Thirdly, the impact of social conditions on reproductive success and offspring performance was different under benign versus harsh thermal conditions for some measures, while in others it was similar. Furthermore, harsh thermal conditions reduced both individual and total parental care. However, sociality had opposite effects: individual care decreased, while total care increased from unifemale-care to biparental-care and multiple-care conditions. To conclude, our study demonstrates that sociality supports greater reproductive success and better offspring performances, but we found no evidence that a more complex sociality during breeding leads to more benefits for both adults and offspring (Moss & While, 2021; Shen et al., 2017). Moreover, we failed to find support for sociality in general shaping the adaptation of organisms to challenging thermal conditions, but we found that the effects of social and ambient thermal conditions on individual reproductive success and performance are not independent.

### 4.1 Effects sociality on reproduction and offspring performance in the two thermal conditions

Our results showed the interactive effects of social conditions and ambient temperatures (*S* × *T_b_*) affected offspring development and performance. Sociality generally buffered the negative effect of harsh temperatures on larval development duration but amplified its impact on dispersed larval weight. Compared to the benign temperature, a shorter duration of larval development was found in the no-care conditions at the harsh temperature, but not in other three social conditions (Fig. 2c). These results might be explained by harsh (i.e. higher) temperatures accelerating carcass decomposition due to increased microbial activity (Conant et al., 2011). However, the presence of parental care mitigates these effects by provisioning food to larvae (Scott, 1998) and suppressing bacterial growth (Shukla et al., 2018), which ascertains enough time for normal development of larvae. The difference in larval mass at dispersal for the biparental-care condition between benign and harsh thermal conditions may be due to males and females employing different strategies in response to thermal variation, affecting offspring performance (Pilakouta et al., 2023). Under both benign and harsh temperatures, there was no difference in larval weight between the unifemale-care and biparental-care conditions, indicating that male presence does not enhance offspring performance. Overall, larvae in the no-care condition were lighter than in conditions with care, suggesting that sociality (i.e., adult care) improves larval weight which is important for short- and long-term offspring fitness (Bartlett, 1987).

We also found a significant effect of the interaction between social conditions and ambient pupating temperatures on offspring lifespan. Interestingly, offspring beetles in the no-care condition exhibited lower mortality rates when pupating under both benign and harsh temperatures, whereas those in the multiple-care condition had higher mortality rates when pupating under benign temperatures. The extended larval development in the no-care condition may give larvae more time to strengthen their immune system. In the yellow mealworm beetle (*Tenebrio molitor*), slow-growing, long-lived individuals have been shown to exhibit enhanced antibacterial activity (Crosland et al., 2024). In contrast, multiple-care condition can potentially impair larval immunity due to high conflict between individuals and alloparental feeding, although concrete evidence is still limited. Furthermore, in the biparental-care condition, newly eclosed beetles that pupated under harsh temperatures faced a higher mortality rate compared to those that pupated under benign conditions. A possible explanation for this result is that, after larval dispersal, larvae and pupae stop feeding and depend on stored energy. Harsh temperatures may increase metabolic rates due to higher energy demands, which could weaken their immune systems (Adamo & Lovett, 2011).

### 4.2 Effects of ambient thermal conditions

Our study shows the adverse effects of harsh ambient thermal conditions on reproductive success and offspring performance. This finding is consistent with previous research that investigated thermal effects on parental care (especially females) and offspring performance in burying beetles (e.g., in *N. vespilloides*, Grew et al., 2019; Pilakouta et al., 2023;2024; in *N. nepalensis*, Malik, et al., 2024; in *N. orbicollis*, Moss & Moore, 2021). The consequences of harsh ambient temperatures for organism reproduction are time-dependent (Pilakouta et al., 2023, 2024), but here we maintained constant temperatures during the breeding phase. Thus, the adverse effects of harsh thermal conditions on reproduction success may have the following reasons. First, high thermal conditions could influence sperm performance and fecundity of adults, subsequently influencing the number of eggs laid and hatching rates (Breedveld et al., 2023; Grew et al., 2019). Unlike birds and some reptiles that incubate their eggs (Deeming, 2001, 2004), burying beetle egg hatching is entirely dependent on soil thermal conditions, which, if suboptimal, can impair hatching (Scott, 1988). These negative effects on egg development may extend to larvae, resulting in smaller brood sizes and lighter larvae at dispersal. Second, harsh thermal conditions have a lasting effect on larval development. High ambient thermal conditions may lead to resource constraints due to exacerbating the decomposition of carcasses (Arce et al., 2012), which subsequently causes intense competition for resources among siblings. Third, the higher ambient temperature may also increase larval metabolic rate and cause thermal damage to the physiological and behavioural activities of individuals (Speakman & Król, 2010), which consequently exhibits a negative effect on brood size and brood mass. Finally, the decreased parental care may also support smaller and lighter broods in harsh thermal conditions. Breeders decreased their parental care under harsh temperatures, which may be explained by (1) balancing the trade-off between reproduction and immune function (Adamo & Lovett, 2011), and (2) saving energy for survival and future reproductive opportunities (Pontzer & McGrosky, 2022). Furthermore, we found that harsh pupating temperatures negatively affected the size of newly-eclosed beetles. This may be due to metabolic damage caused by higher ambient temperatures. Since larvae stop eating after dispersing from the carcass, adult size is largely determined by larval mass (Lock et al., 2004). Adult size influences lifespan, fecundity, and the ability to acquire carcasses for breeding (Bartlett & Ashworth, 1988; Otronen, 1988). Thus, our results suggest that harsh thermal conditions during breeding and pupation significantly impact offspring performance in both the short and long term.

### 4.3 Effects of social conditions

We found that under both thermal conditions the no-care condition resulted in the lowest reproductive success (i.e., brood size and mass) and offspring performance (i.e., average body size of newly eclosed beetles), followed by multiple-care, and then biparental-care and unifemale-care conditions. These findings highlight the significant role of parental care in reproductive advantages (Smiseth et al., 2007; Wang et al., 2021). Also, the presence of parents is found to influence reproduction due to their role in regulating competitive relationships among siblings (Smiseth et al., 2007; Schrader et al., 2015), provisioning developing larvae with predigested carrion (Scott, 1998), and suppressing the spread of microbes (Arce et al., 2012, Shukla et al., 2018), and this is in line with our findings. Moreover, in the no-care condition, there could be strong sibling competition among early hatched, because those larvae obtain the best resources and space for development (contradicting Schrader et al., 2015). Regardless of ambient thermal conditions, there was no difference in reproductive success or offspring performance between unifemale-care and biparental-care conditions, despite differences in individual and total parental care. This suggests that male presence has a limited impact on larval development in this species (Smiseth et al., 2005; Trumbo, 2022). Specifically, (1) males primarily focused on resource guarding rather than directly provisioning larvae (Trumbo, 2022), and (2) females compensated for male absence by increasing their care, leading to minimal effects of the absence of males on reproductive success and offspring performance (Smiseth et al., 2005).

Contrary to our prediction, regardless of thermal conditions, the multiple-care condition resulted in comparable reproductive success and offspring performance to the biparental-care condition but was inferior to the unifemale-care condition, despite significant differences in parental care among these groups. These results suggested that increased coordination (individual variation) in parental care with greater sociality failed to buffer the reproductive costs of harsh breeding temperatures. Possible explanations include: 1) increased conflicts among caregivers and between adults and offspring in the multiple-care condition (Komdeur et al., 2013; Richardson & Smiseth, 2020); 2) hidden collective benefits of multiple-care, such as resource defense, due to limited intra- and interspecific competition under manipulation (Liu et al., 2020; Ma et al., 2022; Tsai et al., 2020); 3) higher risk of pathogen transmission to offspring due to multiple-care (Shukla et al., 2018; Wang & Rozen, 2017). Moreover, greater variation in individual care may enable these individuals to achieve a longer lifespan, thereby pursuing more future reproductive opportunities. However, further comparative and manipulative experiments across animal taxa are needed to test this.

## 5 CONCLUSION

In general, we found that sociality positively influenced reproductive success, offspring performance, and total parental care, but negatively affected larval development duration and individual parental care, while harsh thermal conditions negatively impacted reproductive success, offspring performance, and parental care. Furthermore, a significant interaction between sociality and breeding temperature (*S × T_b_*) affected larval development duration and dispersed larval wight, while the interaction between sociality and pupation temperature (*S × T_p_*) influenced beetle lifespan. These results suggest that sociality buffers the negative effects of harsh temperatures on reproductive success and offspring performance, with limited capacity. Our findings highlight the critical role of sociality in moderating adaptive responses to harsh thermal environments, particularly in the context of global warming.

## ACKNOWLEDGMENTS

This study was supported by a Ph.D. grant from the China Scholarship Council (CSC, 202106180024) to DM, an Ecology Fund of the Royal Netherlands Academy of Arts and Sciences (KNAWWF/807/19021) to LM, and a Dutch Science Council grant (ALW NWO Grant No. ALWOP.531) to JK We thank the Faculty Department of Animal Care, especially Dr. H. Martijn Salomons, for providing frozen mice and lab facilities. We appreciate the constructive comments on the manuscript from Prof. Dr. Jianqiang Li (Beijing Forestry University, China) and Dr. Marco van der Velde (University of Groningen, The Netherlands).

## AUTHOR CONTRIBUTIONS

Donghui Ma designed and conducted the experiment, analyzed the data, and wrote the manuscript. Jan Komdeur and Long Ma supervised the experimental design. Maaike Versteegh oversaw data analysis. LM, JK, and MV edited and revised the manuscript. All authors gave final approval for publication.

## STATEMENT ON INCLUSION

Our study unites authors from two countries, including scientists based in the study’s location. All authors were involved early in the research and design to integrate their diverse perspectives. Regional literature, including relevant work published in the local language, was cited wherever applicable.

## DATA AVAILABILITY STATEMENT

All relevant data will be uploaded to Zenodo.

## SUPPLEMENTARY INFORMATION

**Supplementary Information S1.** Table of Basic information that includes burying beetle breeders, carcass size, and sample size was referred to in our experiment (including the supplementary experiment). All measurements were presented along with means ± standard error.

| Traits | Breeding temperature | No-care | Unifemale-care | Biparental-care | Multiple-care |  |
| --- | --- | --- | --- | --- | --- | --- |
|  |  |  |  |  | Large pair | Small pair |
| Female body size (mm) | 20 °C | 5.06 ± 0.16 | 5.02 ± 0.24 | 5.08 ± 0.15 | 4.91 ± 0.12 | 4.53 ± 0.19 |
| | 23 °C | 5.29 ± 0.25 | 5.18 ± 0.23 | 5.19 ± 0.19 | 5.02 ± 0.07 | 4.61 ± 0.21 |
| Male body size (mm) | 20 °C | 5.10 ± 0.20 | 5.06 ± 0.26 | 5.11 ± 0.18 | 4.94 ± 0.15 | 4.51 ± 0.23 |
| | 23 °C | 5.31 ± 0.29 | 5.21 ± 0.28 | 5.23 ± 0.21 | 5.03 ± 0.10 | 4.68 ± 0.21 |
| Carcass size (g) | 20 °C | 23.51 ± 2.33 | 22.58 ± 2.90 | 22.72 ± 2.96 | 26.32 ± 1.42 | |
| | 23 °C | 21.73 ± 1.31 | 22.04 ± 1.69 | 21.91 ± 1.10 | 25.59 ± 1.23 | |
| Sample size (broods) | 20 °C | 37 | 40 | 39 | 42 |  |
|  | 23 °C | 22 | 25 | 25 | 35 |  |

**Supplementary Information S2**

In accordance with the formal experiment, wherein larvae originating from benign breeding temperatures were exclusively subjected to similar thermal pupation conditions, we conducted a supplementary investigation to assess the impact of harsh thermal conditions during pupation on the fitness of burying beetle offspring reared under benign thermal conditions. Following the outlined experimental design (Figure 1) and procedure, we selected 75 pairs of beetles from the sixth generation of an outbred laboratory population to produce 60 reproductive broods, evenly distributed across four larval-rearing environment treatments. These breeding containers were put into the 20 ℃ climate chamber, with daily monitoring until breeding failure or larvae dispersion occurred. Subsequently, dispersed larvae from each brood were transferred to pupation containers containing moist peat and placed in a climate chamber set to 23 ℃ for pupation. Notably, 11 broods under the ‘no-care’ condition, 13 broods under the ‘unifemale-care’ condition, 14 broods under the ‘biparental-care’ condition, and 14 broods under the ‘multiple-care’ condition produced dispersal larvae successfully. We recorded the breeders’ body size, carcass size, brood mass, and brood size accurately, but not the times of adult beetles being present on the carcass, and the body size of newly eclosed beetles. Consequently, the supplementary experiment data were integrated into select portions of our data analyses, specifically those involving brood mass, brood size, and average larval mass as response variables in our statistical models. Furthermore, we also collected newly eclosed beetles and reared them following our beetles husbandry until death, to record their lifespan.

**Supplementary Information S3.** Table of mean size of newly eclosed beetles and sample size for testing lifespan. All measurements were presented along with means ± standard error, and sample size involving calculation was shown in the brackets. ‘Na’ represents without data.

| Traits | Breeding temperature | Pupating temperature | No-care | Unifemale-care | Biparental-care | Multiple-care |
| --- | --- | --- | --- | --- | --- | --- |
| Size of Newly-eclosed beetles (mm) | 20 °C | 20 °C | 4.79 ± 0.02 (382) | 5.05 ± 0.02 (484) | 4.97 ± 0.03 (456) | 4.85 ± 0.03 (424) |
|  | 20 °C | 23 °C | Na | Na | Na | Na |
| | 23 °C | 20 °C | 4.84 ± 0.04 (122) | 5.08 ± 0.03 (180) | 5.05 ± 0.04 (177) | 4.85 ± 0.04 (235) |
| | 23 °C | 23 °C | 4.63 ± 0.04 (181) | 4.73 ± 0.04 (151) | 4.79 ± 0.04 (175) | 4.61 ± 0.03 (217) |
| Sample size for lifespan (individuals) | 20 °C | 20 °C | 32 | 34 | 28 | 30 |
|  | 20 °C | 23 °C | 29 | 31 | 30 | 32 |
|  | 23 °C | 20 °C | 31 | 29 | 34 | 30 |
|  | 23 °C | 23 °C | 32 | 29 | 30 | 30 |

